# Machine learning analysis of the T cell receptor repertoire identifies sequence features that predict self-reactivity

**DOI:** 10.1101/2022.11.23.517563

**Authors:** Johannes Textor, Franka Buytenhuijs, Dakota Rogers, Ève Mallet Gauthier, Shabaz Sultan, Inge M. N. Wortel, Kathrin Kalies, Anke Fähnrich, René Pagel, Heather J. Melichar, Jürgen Westermann, Judith N. Mandl

**Affiliations:** Data Science group, Institute for Computing and Information Sciences, Radboud University, Nijmegen, The Netherlands; Department of Tumor Immunology, Radboud Institute for Molecular Life Sciences, Radboudumc, Nijmegen, The Netherlands; Department of Physiology and McGill Research Centre on Complex Traits, McGill University, Montreal, Canada; Immunology-Oncology Unit, Maisonneuve-Rosemont Hospital Research Center, Montreal, Canada; Department of Microbiology, Infectious Diseases, and Immunology, Université de Montréal, Montreal, Canada; Institut für Anatomie, Universität zu Lübeck, Lübeck, Germany; Department of Medicine, Université de Montréal, Montreal, Canada

## Abstract

The T cell receptor (TCR) determines the specificity and affinity for both foreign and self-peptides presented by MHC. It is established that self-pMHC reactivity impacts T cell function, but it has been challenging to identify TCR sequence features that predict T cell fate. To discern patterns distinguishing TCRs from naïve CD4^+^ T cells with low versus high self-pMHC reactivity, we used data from 42 mice to train a machine learning (ML) algorithm that predicts self-reactivity directly from TCRβ sequences. This approach revealed that n-nucleotide additions and acidic amino acids weaken self-reactivity. We tested our ML predictions of TCRβ sequence self-reactivity using retrogenic mice. Extrapolating our analyses to independent datasets, we found high predicted self-reactivity for regulatory CD4^+^ T cells and low predicted self-reactivity for T cells responding to chronic infection. Our analyses suggest a potential trade-off between repertoire diversity and self-reactivity intrinsic to the architecture of a TCR repertoire.

## Introduction

During development, each T cell generates one of a possible 10^15^-10^20^ different TCRs through a process of somatic recombination of V and J gene segments for the TCRα chain, and V, D, and J segments for the TCRβ chain.^1–3^ An estimated 90-95% of TCR repertoire diversity is added by terminal deoxynucleotidyl transferase (TdT), a DNA polymerase which mediates non-templated n-nucleotide (nt) additions at the recombining gene segment junctions.^4–8^ The complementarity-determining regions (CDR3) of the TCRα and β chains that span the V(D)J junctions dictate the specificity of a given TCR for peptides presented by MHC molecules (pMHC). The αβCDR3 sequences also determine how strongly each T cell binds to the pMHC ligands it recognizes, whether the peptides being presented derive from self or foreign proteins.^9–11^

That there is a distribution in pMHC reactivity within a T cell population first becomes apparent in the thymus, where the extent of self-reactivity – the average interaction strength with self-pMHC – governs the outcome of positive and negative selection.^12^ Indeed, the self-reactivity of T cells is an important driver of T cell fate even beyond the thymus, not only impacting how well they compete for survival signals in secondary lymphoid organs, but also by establishing pre-wired heterogeneity in gene expression that modulates T cell lineage choice, expansion and memory potential following activation.^13–22^ In addition, both for CD4^+^ and CD8^+^ T cells there is evidence that there is a direct relation between self-reactivity and foreign pMHC binding strength.^14,23^

Despite the pivotal role of the TCR sequence in T cell fate outcomes, few tools exist to systematically define sequence patterns among T cells with similar fates. Substantial progress has been made in developing probabilistic models for the likelihood of generating specific TCR sequences – a key parameter that shapes the pre-selection repertoire and impacts how ‘public’ a TCR sequence is across individuals.^24–27^ For instance, higher intrinsic generation probabilities are thought to explain why TCRs that have a greater proximity to the germline (fewer non-templated nt) are generally more frequent within and across individuals.^28–30^ In addition, recent efforts have made important strides in predicting the foreign antigen specificity of a T cell from its TCR through in-depth studies of epitope-specific TCR repertoires where the T cell ligand is known.^31,32^ Yet, whether it is possible to determine the level of self-pMHC reactivity from the TCR sequence, and hence generate predictions about T cell fate, remains unknown. In part, such studies have been hampered by the very limited numbers of TCR sequences available for which the specific self-peptide recognized has been identified,^33,34^ as well as the technical challenges of measuring the low-affinity binding of self-ligands by T cells.^35^

Here, we make use of a surrogate marker for T cell self-reactivity, CD5, whose surface expression has been shown to vary in direct relation to the strength of sub-threshold tonic self-pMHC signals obtained by T cells.^14,36^ We ask whether there are fundamental differences among TCR sequences from naïve T cells that have distinct self-pMHC binding strengths and whether it might therefore become possible to predict the propensity of a specific T cell to contribute particular effector functions during an immune response. We generated a dataset of 1.5×10^7^ unique CDR3β sequences from a total of 42 mice, investigating patterns among TCRβ chain sequences between mature CD5^lo^ and CD5^hi^ naïve CD4^+^ T cells, as well as sequences in the double positive (DP, pre-selection) and single positive (SP, post-selection) stage in the thymus.

The vast majority of the TCRβ sequences we observed both within and between mice appeared only once (i.e., were entirely private); this necessitated the development of novel analysis tools to compare sequences between sorted T cell and thymocyte populations. Implementing a machine learning (ML) algorithm, we showed that it was possible to identify subsets of TCRβ sequences that were strongly associated with low or high self-reactivity, despite a large overlap between sequences from sorted T cell populations. Our analyses revealed that CD5^hi^ TCRβ sequences are more germline-like (fewer non-templated nt additions) and have specific features at the amino acid (aa)-level. Moreover, CDR3β sequences from T cells with high self-reactivity are enriched among CD4^+^ SP thymocytes compared to DP thymocytes, suggesting they are more rapidly positively selected. Through experimental validation using TCR retrogenic mice with TCRβ sequences not in our original dataset, as well as analysis of publicly available datasets, we made the surprising finding that even without a known TCRα sequence, the TCRβ sequence can provide information on the relative self-reactivity of naïve CD4^+^ T cells and shed light on repertoire differences in other contexts, such as acute versus chronic infection. Overall, our use of an ML model to stratify complex TCR sequencing datasets provides fundamental insight into the architecture of TCR repertoires.

## Results

### *Dntt* expression correlates with CD5 levels on CD4^+^ T cells suggesting a possible impact on TCR sequence

For both naïve CD4^+^ and CD8^+^ T cell subsets, gene expression comparisons unexpectedly identified *Dntt* as one of the most enriched genes in T cells with low self-pMHC reactivity.^17,21,37,38^ Since *Dntt* is the gene that encodes for the non-templated DNA polymerase TdT, this suggested a possible relation between a T cell’s TCR sequence and its self-reactivity. To investigate the relation between the self-reactivity of CD4^+^ T cells and the expression level of *Dntt* in more detail, we first asked whether the *Dntt* expression difference was due to residual mRNA present among recent thymic emigrants (RTE). In line with a heightened sensitivity to pMHC engagement by RTE,^39^ GFP^+^ RTE identified in Rag-GFP reporter mice were enriched among CD5^hi^ cells, rather than among CD5^lo^ cells **(Supplementary Figure S1A)**. Nevertheless, we excluded GFP^+^ RTE cells when sorting on naïve CD4^+^ T cells expressing low, mid, and high levels of CD5 to determine *Dntt* mRNA expression by qPCR for each sorted population. Consistent with prior data, we found an inverse relation between surface CD5 levels and *Dntt* expression by naïve CD4^+^ T cells, with CD5^lo^ cells expressing 15-fold more *Dntt* transcripts than CD5^hi^ cells **(Figure 1A)**. Of note, in CD5^lo^ cells, *Dntt* expression was substantially lower (∼ 40-fold) than in double negative (DN) and DP thymocytes that are in the process of actively rearranging their TCRα and β chains, respectively (**Figure 1A**). No TdT protein was detected in mature naïve T cells either (**Supplementary Figure S1B**). Together, these data led us to hypothesize that differing levels of *Dntt* expression during thymic development between individual T cells might play a role in determining the number of non-templated nt added to the TCR V(D)J junctions, and therefore contribute to determining the self-pMHC reactivity of a T cell.

**Figure 1:**
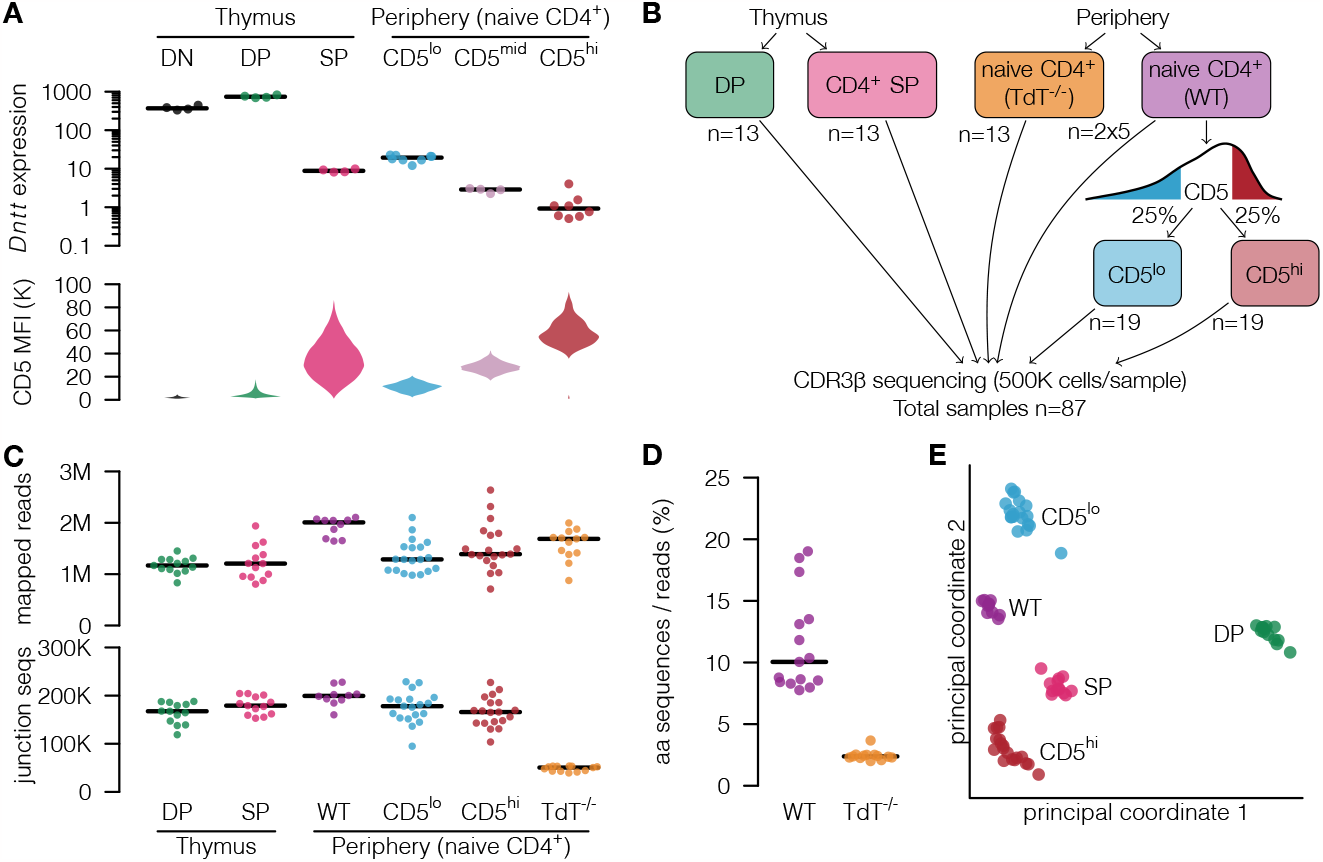
Study design and sample characteristics to investigate the relation between TdT expression and self-reactivity in CD4^+^ T cells. **(A)** CD5 protein and *Dntt* mRNA expression during thymic development and across peripheral naïve CD4^+^ T cells sorted into 20% lowest, mid and highest CD5-expressing populations (CD5^lo^, CD5^mid^ and CD5^hi^, respectively). DN, double negative thymocytes; DP, pre-selection double positive thymocytes; SP, CD4^+^ single positive thymocytes. **(B)** Schematic of samples used. The DP (CD4^+^ CD8^+^ TCRβ^lo^ CD69^-^) and SP (CD8^-^ CD25^-^ TCRβ^+^ CD3^+^) thymocytes, 25% CD5^lo^ and CD5^hi^ naïve CD4^+^ T cells (CD44^lo^ CD62L^+^ CD25^-^ TCRβ^+^ CD4^+^) were sorted from 13 mice, with 6 additional mice included for the CD5^lo^ and CD5^hi^ populations (N=19). Total naïve CD4^+^ T cells were sorted from 13 TdT^-/-^ mice, and two sets of total naïve CD4^+^ T cells samples were sorted from 5 wild type (WT) mice (labelled as 2×5). For each population, the CDR3β region of the TCR was sequenced for a total of 500,000 cells. **(C)** Number of mapped reads and unique CDR3β amino acid (aa) sequences per sample. **(D)** Diversity in CDR3β aa sequences identified in naïve CD4^+^ T cells from TdT^-/-^ compared to WT mice. **(E)** Multidimensional scaling overview of cell populations sorted from WT mice, based on the overlap (Jaccard index) in junction aa sequence, V region, and J region.

To comprehensively characterize the TCR sequences present in a naïve CD4^+^ T cell population and relate this to self-reactivity, as well as thymic selection, we performed deep sequencing of the TCR CDR3β regions. Per sample, we sorted 5×10^5^ total CD4^+^ T cells, as well as CD5^lo^ and CD5^hi^ naïve CD4^+^ T cells (top and bottom 25% of the CD5 distribution) from wild type (WT) mice, and as a comparison TCR repertoire, naïve CD4^+^ T cells from TdT^-/-^ mice (**Figure 1B** and **Supplementary Figure S1C**,**D**). Notably, our prior work showed that this sorting strategy excludes all regulatory (FoxP3^+^) CD4^+^ T cells, which express more CD5 on average.^21^ To investigate the representation of TCRβ sequences through thymic development, we sorted on DP thymocytes that had not yet received a TCR signal and SP CD4^+^ thymocytes that had been positively selected but not yet left the thymus from the same mice (**Figure 1B** and **Supplementary Figure S1E**,**F**). We successfully mapped ∼1.3 million reads in each WT sample to the CDR3β region and found on average 150K unique aa sequences (**Figure 1C**). For TdT^-/-^ samples, we mapped similar numbers of sequences per sample (∼1.6 million) but, as expected, the number of unique aa sequences was greatly reduced (**Figure 1C**), corresponding to a 73% reduction in median estimated diversity (**Figure 1C**,**D**). Overall, basic sample statistics clearly differed between WT and TdT^-/-^ naïve CD4^+^ T cells but not between the different WT subpopulations (**Figure 1C**). We next investigated the degree of overlap in TCR sequences between the WT samples as quantified by the Jaccard index. For this analysis, and throughout this paper unless stated otherwise, we considered two CDR3β sequences to be identical if they had the same mapped V gene, the same mapped J gene, and the same junction sequence (including the remainder of the D gene) at the aa-level. A multi-dimensional scaling plot of a distance matrix based on the Jaccard index (**Figure 1E**) showed a clear separation between the samples according to their phenotypes.

### Large overlap and few differences in the CDR3β repertoires of CD5^lo^ and CD5^hi^ naïve CD4^+^ T cells

Our multi-dimensional scaling analysis (**Figure 1E**) showed that two repertoires from the same T cell population in distinct mice shared a greater number of TCR sequences than two repertoires from different T cell populations obtained from the same mouse, suggesting the existence of T cell population-specific features in their TCR sequences. To further investigate the magnitude of TCR sequence overlap, we first quantified the number of mice each sequence was observed in. Almost 80% of the CD5^lo^ and CD5^hi^ sequences occurred only in a single mouse (private sequences), while less than 5% were seen in 5 mice or more (**Figure 2A**,**B**). These results indicated a very low overlap between the TCR repertoires of sorted T cell populations, but because we sequenced only a fraction of the full CDR3β repertoire, we were likely underestimating the true overlap. To account for this and calculate the expected overlap between repertoires given our sequencing setup, we also sequenced duplicate sets of naïve CD4^+^ T cell samples from the same mouse (**Figure 2C**). The overlap between the pairs of naïve CD4^+^ T cell repertoires taken from the same mouse was ∼9-10%, which was not much greater than the overlap between CD5^lo^ and CD5^hi^ repertoires at ∼6-8% (**Figure 2D**). Thus, normalizing by the duplicate sample sets showed that TCR sequences among CD5^lo^ and CD5^hi^ naive CD4^+^ T cells likely overlapped substantially.

**Figure 2:**
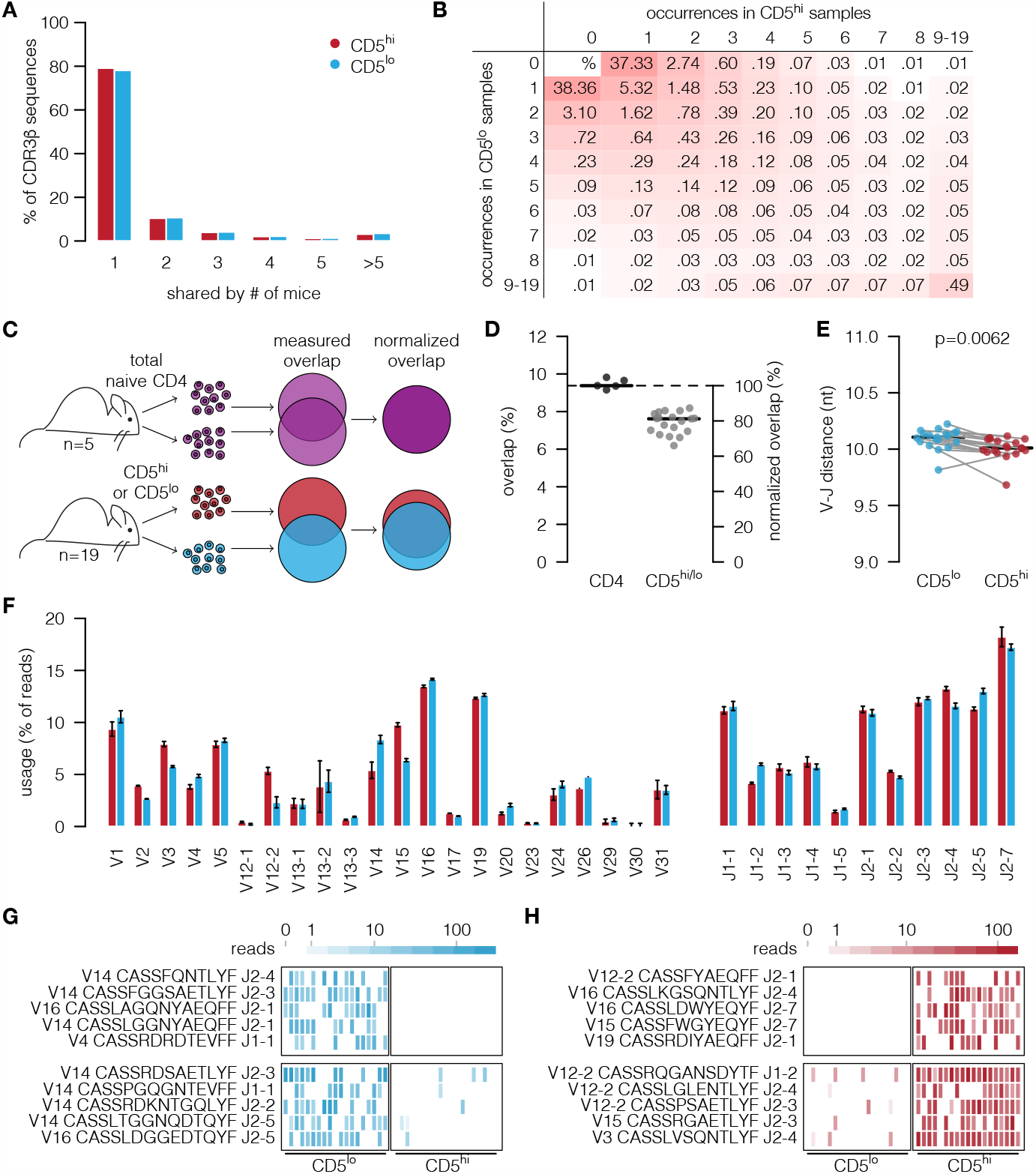
Naïve TCR repertoires are largely private and there is extensive overlap within mice between CD5^lo^ and CD5^hi^ populations. **(A)** Sharing distribution of CDR3β aa sequences from CD5^lo^ and CD5^hi^ naïve CD4^+^ T cells across 19 mice. **(B)** Table of occurrence patterns indicating percent of CDR3β aa sequences found across mice and between sorted CD5^lo^ and CD5^hi^ populations. **(C**,**D)** Duplicate samples from individual mice of WT total naïve CD4^+^ T cells were used to estimate the maximal within-mouse overlap expected (C), and thus estimate the actual sequence overlap (D) in CDR3β aa sequences from CD5^lo^ and CD5^hi^ naïve CD4^+^ T cells sampled from the same mouse. **(E**,**F)** V-J distance in nt (E), and V/J gene segment usage (F), in CD5^lo^ and CD5^hi^ naïve CD4^+^ T cells. Error bars: bootstrapped 95% confidence interval of the estimated proportion (N=19). **(G**,**H)** Top row: Top 5 most differentially expressed CDR3β sequences found only in CD5^lo^ samples (G) or CD5^hi^ samples (H). Bottom row: Top 5 most differentially expressed CDR3β sequences found in both CD5^lo^ samples and CD5^hi^ samples (H).

With this caveat in mind, we next asked whether there were systematic differences between TCR sequences from CD5^lo^ and CD5^hi^ CD4^+^ T cells. We tested the hypothesis that thymocytes expressing higher levels of TdT preferentially gave rise to CD5^lo^ cells with longer CDR3β junction sequences (which then retained relatively higher expression of *Dntt* as mature naïve T cells even though the gene was largely silenced)(**Figure 1A**). While there was a robust difference in nt length between CD5^lo^ and CD5^hi^ cells, the difference was small (**Figure 2E**), and most differences in V and J gene segment usage remained well below 20% (**Figure 2F**). Despite the absence of easily detectable sequence features that strongly differed between the TCR sequence data from CD5^lo^ and CD5^hi^ cells, a standard differential gene expression (DGE) analysis found 1131 differentially expressed sequences (FDR < 0.05), some of which were exclusively found in either CD5^lo^ and CD5^hi^ cells (total of 767 sequences) or substantially enriched in one population over the other (examples of both are shown in **Figure 2G**,**H**). While it was encouraging that we could detect sequences that were specifically enriched in either CD5^lo^ or CD5^hi^ cells, a set of only ∼1000 sequences (out of a total of ∼3.5 million) provided a very limited number from which to discern patterns with regard to self-reactivity. Moreover, conventional DGE analyses almost exclusively identify sequences that are public (appear across several mice to achieve statistical significance), which likely introduces specific sequence biases as it has already been described that public sequences have reduced n-nucleotide additions^28,40^ and tend to contain the more common V and J gene fragments. Thus, we went on to develop an alternative approach to identify predictive features associated with high or low self-reactivity from the full dataset, including the private sequences.

### An ML algorithm can distinguish between the CDR3β repertoires of CD5^lo^ and CD5^hi^ cells

The large overlap we found between the repertoires of CD5^lo^ and CD5^hi^ naïve CD4^+^ T cell populations (**Figure 2D**) indicated that for many CD4^+^ T cells the CDR3β sequence alone does not determine the CD5 expression level. This could be due to a number of reasons, including: (1) the TCRα chain may play an important role in the self-reactivity of a T cell; (2) the sets of self-peptides that different T cells with the same TCR encounter during thymic selection might be different, modulating CD5 expression; and/or (3) there might be large stochasticity in TCR signal strength obtained from interactions with self-peptides in the thymus, depending on, for instance, the intervals between encounters. To investigate whether, despite the overlap in repertoires, there were CDR3β sequence features that distinguished TCRs from T cells with high compared to low self-reactivity and could be identified directly from raw sequences, we turned to an ML algorithm.

The ML algorithm we implemented was a binary classifier constructed to distinguish two different classes of CDR3β sequences from each other. The general procedure of constructing and training this classifier is independent of the specific classes being distinguished (such as CD5^lo^ versus CD5^hi^), and we designed it to ignore the frequency at which TCR sequences occur in one sample or across samples, such that the classifier devotes equal attention to “private” and “public” sequences. Specifically, we divided the data into a training dataset and a testing dataset, setting aside 20% of the TCRβ sequences from each of the two T cell populations of interest for later testing. We removed from the testing dataset all sequences that also occurred in the training set; specifically, we removed all sequences from the testing set if they shared the same V gene, J gene, and aa junction sequence with a training sequence. For the training dataset, we removed all TCR sequences that were found in both populations. Here we considered two sequences the same if they shared the same V gene, J gene, and nucleotide junction sequence, as we expected the number of n-nucleotides to be of interest, which cannot be discerned from the aa sequence. Finally, we balanced the training data by removing sequences from the larger population at random to obtain two equally large training sets for each class, avoiding potential bias for the larger class (**Figure 3A**). Notably, we used aa-based sequence equality for constructing our test sets because this ensured that the network did not “cheat” and score well on the test data by simply learning the translation rules from nt to aa.

**Figure 3:**
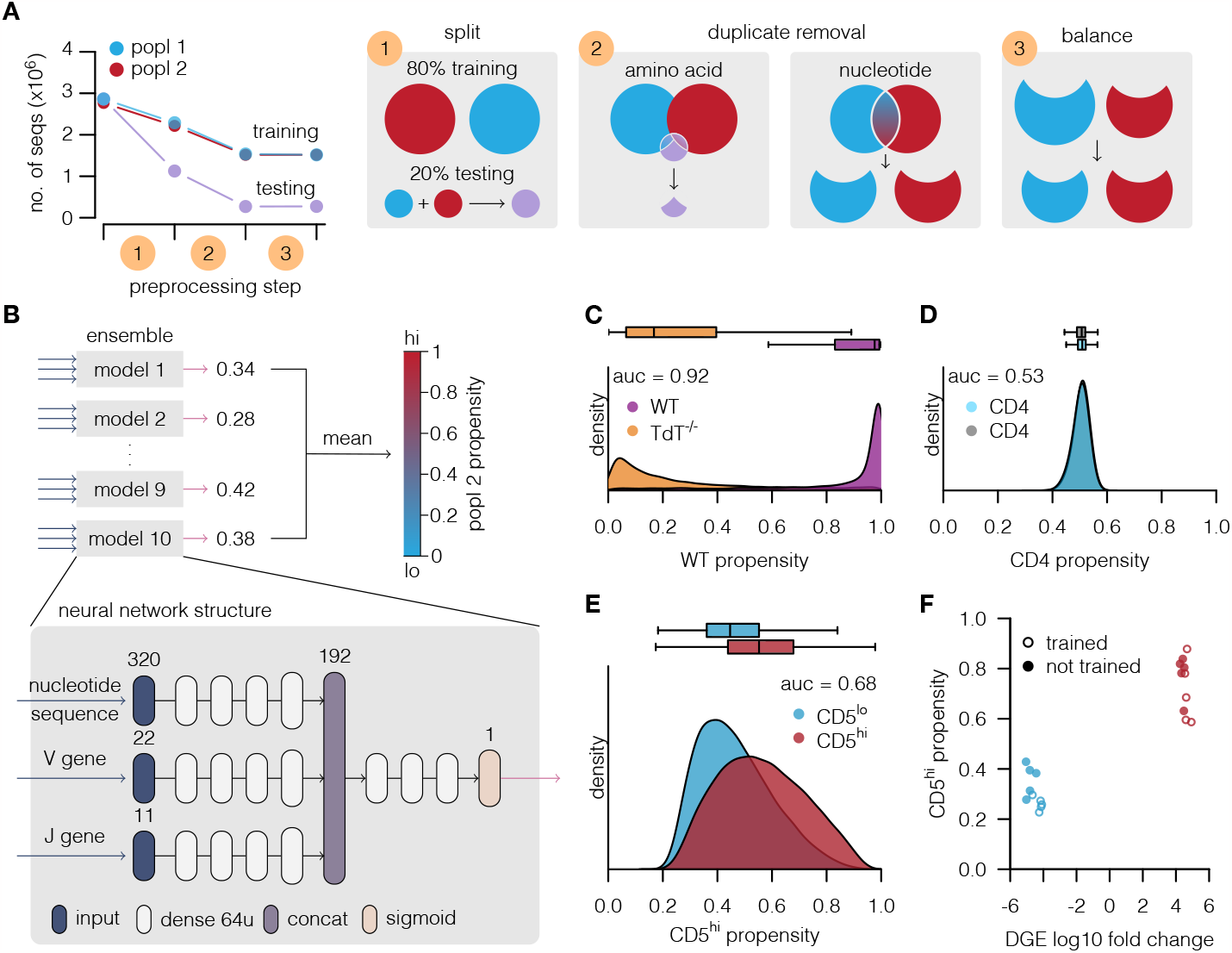
Machine learning can distinguish between TCRβ sequences from CD5^lo^ and CD5^hi^ CD4^+^ T cells. **(A)** Schematic of our training setup; sequence numbers given in leftmost panel are for CD5^lo^ versus CD5^hi^ classification. We set aside 20% of the sequences for testing the model performance; panels C-E are based on these test sets. To avoid information sharing between training and test data, we remove all CDR3β sequences that map to the same aa sequence from the training data. Finally, we balance the training data using sub-sampling to avoid bias towards either population. **(B)** Schematic of the artificial neural network (ANN) architecture used to distinguish between TCR from two populations (popln 1 and 2). Each ANN is given an input TCR sequence, V and J gene segment usage, and outputs a number between 0 and 1 (propensity), where 1 is certainty that the TCR is from popln 2, and 0 is certainty that the TCR is from popln 1. We average such predictions over an ensemble of 10 ANNs. **(C)** ML-determined WT propensity distributions, comparing naïve CD4^+^ T cell TCRβs from WT and TdT^-/-^ mice. Auc, area under curve, assesses overlap of distributions (auc=0.5, complete overlap; auc=1, no overlap). **(D)** ML-determined CD4 propensity score distributions, comparing naïve CD4^+^ T cell TCRβs sequenced from duplicate samples taken from the same mice. **(E)** ML-determined CD5^hi^ propensity score distributions, comparing TCRβs from CD5^lo^ and CD5^hi^ naïve CD4^+^ T cells. **(F)** ML-determined mean CD5^hi^ propensity scores for the top 10 most enriched TCRs sequences present in CD5^lo^ versus CD5^hi^ naïve CD4^+^ T cells identified in **Figure 2G**,**H**.

The ML algorithm we used was a simple feed-forward neural network with seven hidden layers (**Figure 3B**); when experimenting, we found this network structure to perform substantially better than an even simpler one-layer network, but we did not observe substantial additional performance gains when adding more complexity (data not shown). We followed the CDR3β input encoding previously proposed^41^ and provided the ML algorithm with the mapped V and J genes and the CDR3β junction nt sequence as input. For each individual TCRβ sequence, the network output was a number between 0 and 1 indicating how likely it deemed the sequence to belong to the (arbitrary) reference class. We refer to this number as a propensity score. For example, when training a network to distinguish CD5^lo^ and CD5^hi^ sequences and using CD5^hi^ as a reference class, we refer to the resulting number as the CD5^hi^ propensity score. Since the process of training a neural network is stochastic, and repetition of the training can change the prediction of the network, we worked with an ensemble of 10 networks to reduce this stochasticity, taking the mean propensity score of the 10 outputs, thus also being able to evaluate the reproducibility of the ML predictions (**Figure 3B**).

To investigate the performance of the ML algorithm, we performed two controls. First, we showed that we were readily able to distinguish WT from TdT^-/-^ TCR repertoires (**Figure 3C**; area under the receiver operating curve [auc] value: 0.92, with 1 indicating perfect classification and 0.5 being a random coin toss). Second, we confirmed that we were unable to distinguish TCRβ repertoires from the two duplicate sets of sorted naïve CD4^+^ T cell populations (**Figure 3D**), indicating that the ML algorithm was not picking up spurious or unexpected patterns in our TCR sequencing datasets. Interestingly, the ML classifier was able to distinguish between CD5^hi^ and CD5^lo^ TCR sequences on the population level (**Figure 3E**). While many CDR3β sequences had propensity scores around 0.5, indicating no confident predictions could be made, a clear shift in the propensity distributions was nevertheless visible (auc=0.68) in comparing the CD5^hi^ and CD5^lo^ naïve CD4^+^ T cell populations. Moreover, evaluation of the differentially expressed sequences we identified earlier (**Figure 2G**,**H**) showed a clear alignment between our ML propensity scores and the DGE analysis, regardless of whether or not the DGE sequence was in the training set (**Figure 3F**).

Thus, a relatively simple ML algorithm was able to discriminate CD5^lo^ and CD5^hi^ TCRβ repertoires on the whole-repertoire level, even though many individual TCRβ sequences could not be confidently assigned either way. Therefore, we next investigated which specific sequence patterns were enabling the ML-based discrimination.

### Characterizing confidently ML-classified CDR3β sequences

To establish TCRβ sequence features of T cells with high or low self-reactivity, we next used the ML algorithm as a filter to identify cells with confidently classified CD5 status (high or low propensity scores) for further analysis. To do so, we ranked each CD5^lo^ and CD5^hi^ sample by the ML-assigned CD5^hi^ propensity score, and then extracted the bottom or top 15% of the CD5^lo^ and CD5^hi^ sequences, respectively, for further analysis (**Figure 4A**). We denoted these selected TCRβ sequences as “confidently predicted” CD5^lo^ and CD5^hi^ (coCD5^lo^ or coCD5^hi^, respectively). First, to revisit our hypothesis of the differential role for TdT in generating TCRs with high compared to low self-reactivity, we compared β- chain VDJ junction lengths between coCD5^lo^ and coCD5^hi^ sequences and found a greater difference than was observed for the unfiltered CD5^lo^ and CD5^hi^ sequence datasets (compare **Figure 2E** with **Figure 4B**). Since the 0.5nt length difference between the confidently predicted CD5^lo^ and CD5^hi^ sequences still appeared to be small, we aimed to put this in perspective and estimated the number of non-templated nt directly by mapping the D segment to the junction and counting the nt that could not be explained by this mapping. This analysis showed that coCD5^lo^ sequences had ∼15% more non-templated nt than coCD5^hi^ sequences (**Figure 4C**). Together, analyses of the ML-filtered CD5^lo^ and CD5^hi^ TCR sequence datasets suggested that there is indeed a pattern with regard to strength of self-reactivity and TdT-dependence of a given TCRβ sequence, as hypothesized.

**Figure 4:**
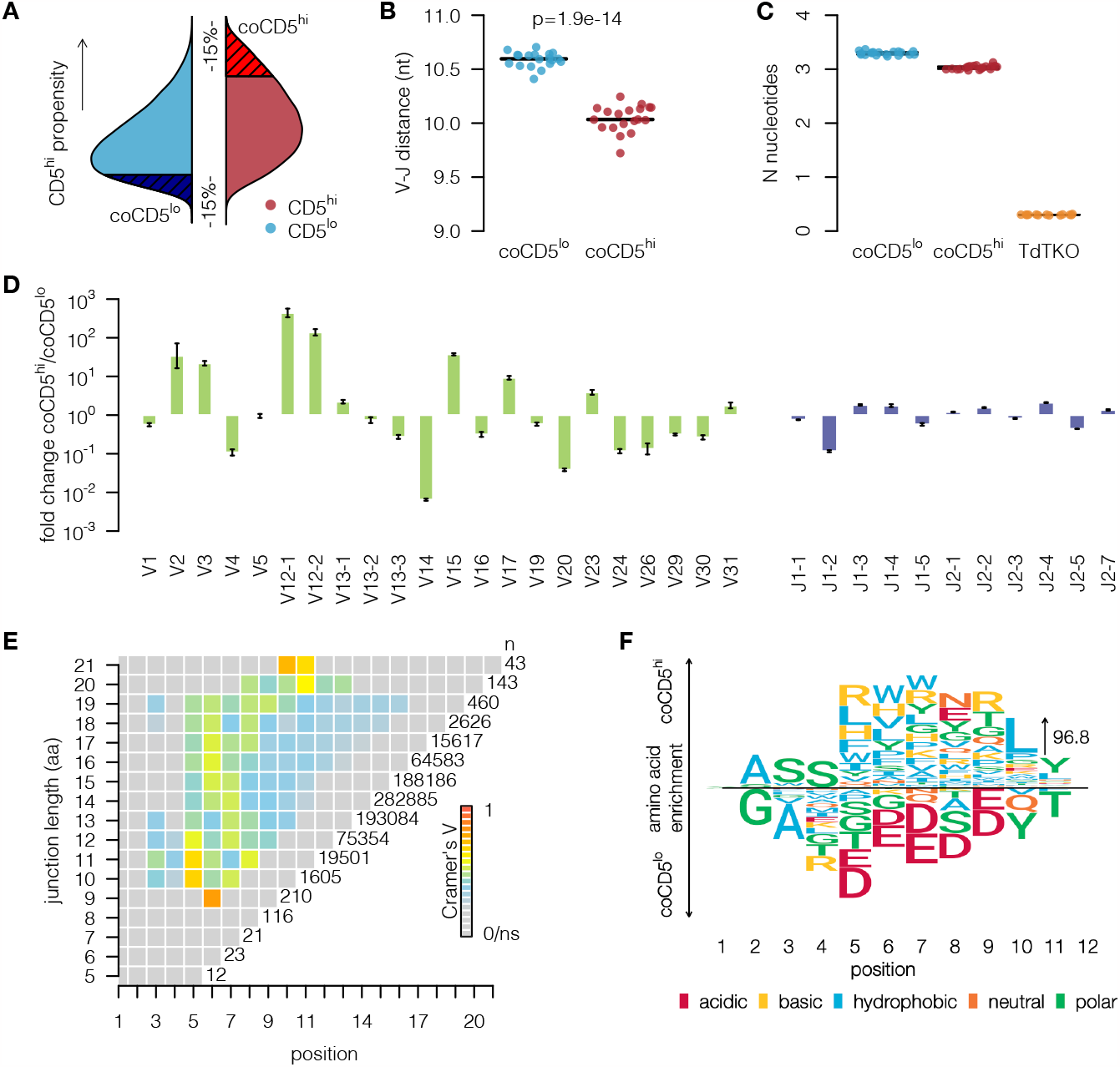
CDR3β sequences with confident ML CD5^hi^ propensity scores have distinct aa usage, number of N-nucleotide additions, and junction length. **(A)** ML-scored CDR3β sequences falling into the bottom (coCD5^lo^) or top (coCD5^hi^) 15% of the CD5^hi^ propensity score distribution were selected for further analysis. **(B**,**C)** V-J distance (B), and estimated number of n-nucleotide additions (C) for coCD5^lo^ and coCD5^hi^ TCRβ sequences, compared to TdT^-/-^ sequences for reference. **(D)** Fold difference between V/J usage of coCD5^lo^ and coCD5^hi^ CDR3β sequences. Error bars: Bootstrapped 95% confidence intervals. **(E)** Differences between observed and expected aa distributions quantified using Cramér’s V for each position in the CDR3β sequence, indicating specific aa preference patterns in the middle positions. TCRβ sequences are stratified by junction length, with number of sequences per length (n) shown to the right of each junction length. **(F)** Amino acid preference patterns for CDR3β sequences with length 12 (from E). Amino acids more frequent in coCD5^lo^ are below the line and in coCD5^hi^ are above the line. Arrow height indicates an enrichment of aa frequency that is 96.8-fold larger than expected by chance.

Next we asked whether there were other features of TCRβ chains identified as coCD5^lo^ and coCD5^hi^ by the ML algorithm. In examining V gene segment usage, we found large (in some instances exceeding 100 fold) differences between coCD5^hi^ and coCD5^lo^ sequences, with, for instance, V14 and V20 gene segments enriched among coCD5^lo^ sequences, while V12-1, V12-2 and V15 gene segments were greatly enriched among coCD5^hi^ sequences (**Figure 4D**). Notably, these V gene segment usages were also represented in the differentially expressed TCRs identified earlier (**Figure 2G**,**H**), but were not visible in the full dataset without applying the ML filter (**Figure 2F**). While more muted, we also observed differences in J gene segment usage, including a ∼10-fold over-representation of J1-2 among coCD5^lo^ sequences (**Figure 4D**). Interestingly, when we stratified the ML-identified coCD5^lo^ and coCD5^hi^ sequences by length, we were able to pinpoint specific aa positions in the CDR3β junction sequence that differed significantly between confidently predicted high and low CD5-expressing T cells. The largest divergence in coCD5^lo^ and coCD5^hi^ TCRs was generally found near the middle of the TCRβ sequence at positions 5-7 (**Figure 4E**). Further, examining the probability of individual aa appearing in each CDR3β position, we noted that there were clear patterns. For instance, coCD5^lo^ cells were consistently enriched for aspartic and glutamic acid (both acidic, negatively charged aa), while coCD5^hi^ sequences were enriched for the hydrophobic aa leucine, tryptophan, valine, and phenylalanine, as well as basic (positively charged) aa, arginine, lysine and histidine (**Figure 4F**).

Taken together, our use of the ML algorithm to filter on CDR3β sequences based on propensity scores allowed us to better understand and characterize the differences between TCRβ sequences represented in CD5^lo^ compared to CD5^hi^ naïve CD4^+^ T cells. Through these analyses we found that there are specific patterns regarding VDJ junction length, numbers of n-nt added, V and J segment usage, as well as characteristics of aa represented particularly in positions 5-7 of the TCRs in our dataset that were predictive of T cell self-reactivity.

### ML-derived TCRβ sequence features predict self-reactivity in a validation dataset

We next wanted to establish to what extent the TCRβ sequence features identified by our analysis of confidently ML-classified sequences were able to predict CD4^+^ T cell self-reactivity *without* the use of an ML algorithm. We therefore trained a logistic regression model – a simple statistical classifier – to distinguish CD5^lo^ and CD5^hi^ CDR3β sequences based on 11 features: V-J distance, usage of acidic and hydrophobic amino acids, usage of the V genes 2, 3, 12-1, 12-2, 14, 15, 20, and usage of J1-2. The logistic regression model was able to distinguish CD5^lo^ and CD5^hi^ CDR3β sequences (**Figure 5A**) when trained and evaluated on the same data as our ML model (**Figure 3A**), although it discriminated the sequences less well (auc=0.58) than our full ML model. This finding supported the hypothesis that these features contribute to predicting self-reactivity.

**Figure 5:**
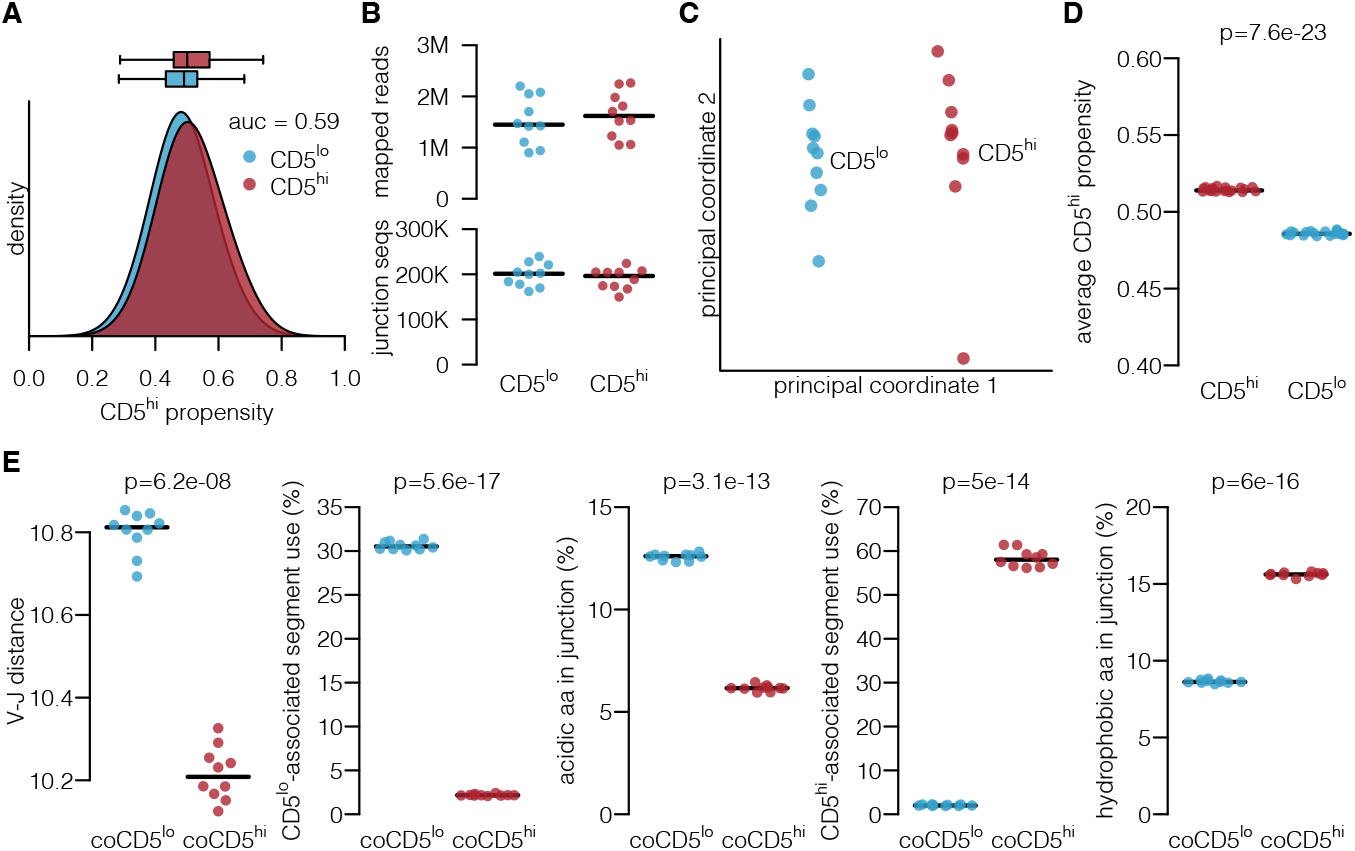
ML-derived features predict CD4^+^ T cell self-reactivity in an independent dataset. **(A)** Performance of a logistic regression model consisting of 11 features when trained and evaluated on the same data as our ML model (**Figure 3E**). **(B)** Mapped reads and unique junction sequences in a validation dataset sequences from 10 mice. **(C)** Multidimensional scaling plot of the validation dataset, as in **Figure 1E. (D)** Average CD5^hi^ propensity score from the logistic regression model for each sample, compared between CD5^lo^ and CD5^hi^ T cells. P-value: two-tailed paired T test. **(E)** Average values of ML-derived features (V-J distance, acidity, hydrophobicity) or groups of features (CD5^lo^ associated segments: V14, V20, or J1-2, CD5^hi^ associated segments: V2, V3, V12-1, V12-2, or V15) for ML-filtered validation sequences (coCD5^lo^ and coCD5^hi^, see **Figure 4A**). P-values: two-tailed paired T tests, Bonferroni adjusted.

To provide a more stringent test of our sequence feature predictions, we collected an independent dataset consisting of CD5^lo^ and CD5^hi^ CDR3βs sequenced from an additional ten mice (**Figure 5B**,**C**). We used this validation dataset to perform pre-planned analyses, publishing the analysis plan ahead of its execution.^42^ First, we used the logistic regression model that was trained on our previously acquired data to determine the average CD5^hi^ propensity of all sequences in each dataset without using the ML algorithm as a filter. We found that the obtained CD5^hi^ propensity scores were substantially higher for CD5^hi^ CDR3β sequences than for those from CD5^lo^ sorted samples, suggesting that even in an independent dataset we were able to predict CD5 status based on the ML-determined TCR sequence features (**Figure 5D**). Further, when examining the average value of each TCR sequence feature (grouping V and J gene usage by CD5^lo^- and CD5^hi^-association for simplicity) in ML-filtered sequence subsets (i.e., coCD5^lo^ and coCD5^hi^ samples determined from the validation data), we found that each of them differed with CD5 status (**Figure 5E**).

In summary, our validation analyses conducted on an additional dataset showed that the ML-identified sequence features on their own were able to predict self reactivity.

### CD5^hi^ TCRβ sequences are more efficiently positively selected in the thymus

The level of CD5 on the surface of naïve T cells is determined during thymic development, although it can be modulated in the periphery by access to self-pMHC.^21,43^ Given the skewed CD5 distribution of naïve CD4^+^ T cells, with a greater number of T cells being CD5^hi^, it has been postulated that T cells with greater self-pMHC reactivity are more likely to be selected in the thymus.^10,14^ Indeed, this hypothesis would be consistent with data suggesting that T cells from TdT^-/-^ mice are more efficiently positively selected and thus that the germline-encoded T cell repertoire is inherently more self-reactive.^44^ However, whether greater self-reactivity leads to increased positive selection efficiency has not been examined at the TCR sequence level. Therefore, we next sought to compare the CDR3β repertoires of thymocytes at the pre-selection DP and post-selection SP thymic selection stages to those of CD5^lo^ and CD5^hi^ naïve CD4^+^ T cells and ask whether sequence biases could be detected that would suggest that some TCRs are more efficiently selected than others.

To do so, we used the same ML architecture as before (**Figure 3B**) to train a classifier to distinguish the TCRβs from the set of sorted DP from those of SP thymocytes sequenced from the same mice (N=13; **Figure 1B**). We found that the ML algorithm was able to detect differences between DP and SP TCRβ sequences, with a clear shift in SP propensity scores between the two thymocyte populations in a separate test set (auc = 0.7, **Figure 6A**). Importantly, all SP TCR sequences must have previously passed through the DP stage, and correspondingly, our classifier did not achieve high propensity scores (>0.8) for many SP thymocyte sequences. Conversely, consistent with there being DP TCR sequences that rarely reach the SP stage, our ML classifier assigned very low scores (<0.2) for a subset of sequences from DP thymocytes (**Figure 6A**). Next, as we did for the CD5^lo^ and CD5^hi^ TCR sequence comparisons, we focused further analysis on the confidently predicted coDP and coSP TCRβ sequences (bottom and top 15% of the SP propensity scores). We found that DP CDR3βs with very low SP propensity scores were significantly longer (V-J distance difference of ∼5 nt) than coSP TCRs (**Figure 6B**), suggesting that a subset of the coDP CDR3β sequences were too far removed from the germline to be positively selected. Analysis of aa usage patterns showed that acidic (negatively charged) aa were enriched in the coDP sequences, similarly to what we found in CD5^lo^ naïve CD4^+^ T cells (**Figure 4F**), and there was also evidence for enrichment in hydrophobic, polar (glycine, threonine and tyrosine), and non-charged (asparagine and glutamine) aa in coDP compared to coSP CDR3β (**Figure 6C**).

**Figure 6:**
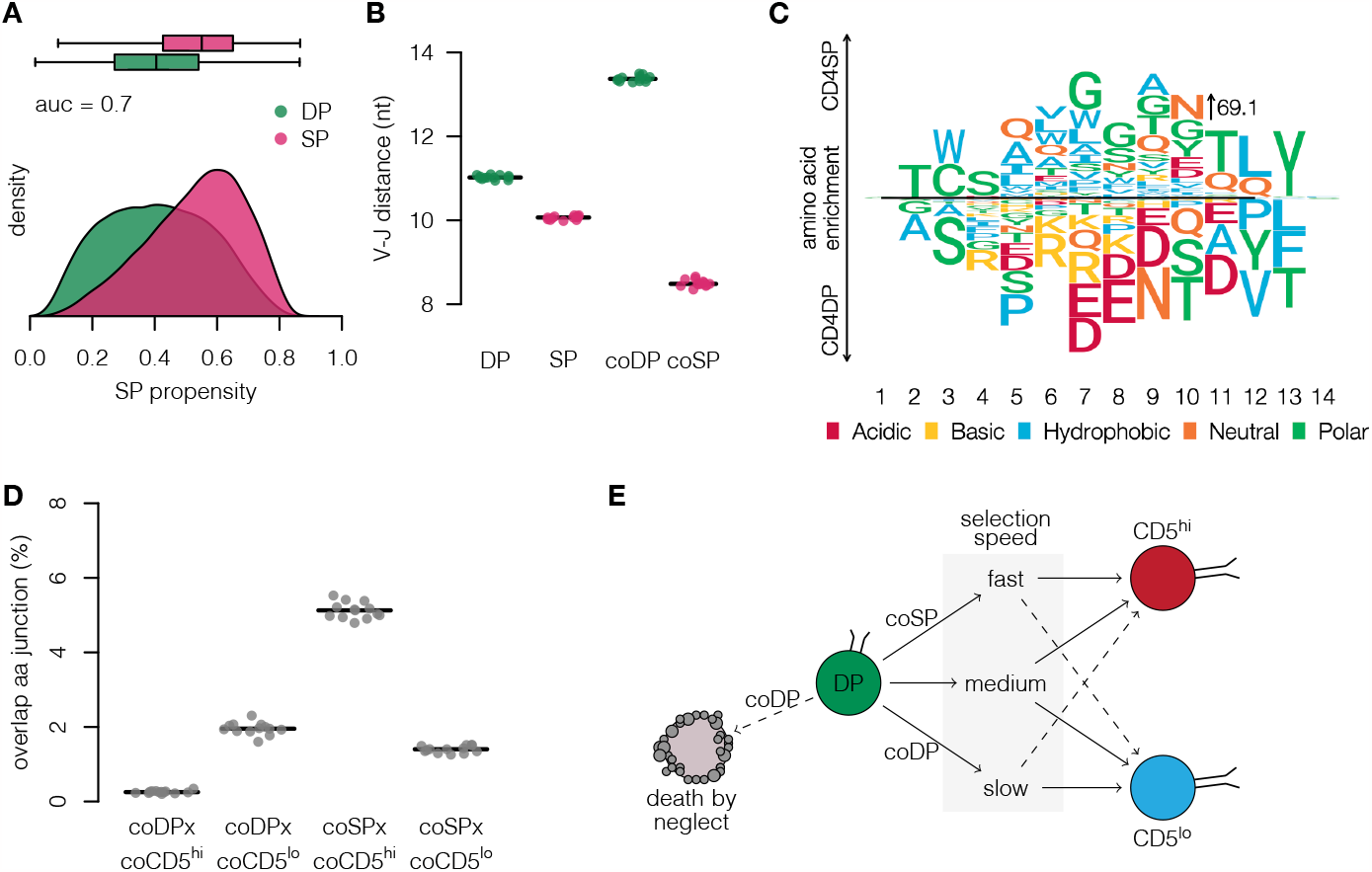
Machine learning identifies subsets of TCRβ sequences with distinct thymic selection fates based on their self-reactivity. **(A)** ML-determined single-positive (SP) propensity score distributions, comparing TCRβs from pre-selection DP and CD4^+^ SP thymocytes. **(B)** Junction length for all DP and SP, as well as ML-determined coDP and coSP TCRβ sequences. **(C)** Amino acid preference patterns for CDR3β sequences with length 12. Amino acids more frequent in coDP are below the line and in coSP are above the line. Arrow height indicates an enrichment of aa frequency that is 69.1-fold larger than expected by chance. **(D)** Overlap of CDR3β sequences between coDP and either coCD5^lo^ or coCD5^hi^, or between coSP and either coCD5^lo^ or coCD5^hi^ TCRβ sequences. **(E)** Schematic model for explaining the findings in A-D. coDP sequences represent those that rarely become SP while coSP sequences are those that rapidly become SP. Rapid selection should correlate with higher propensity to become CD5^hi^, while remaining in the DP stage for a long time may reflect lower binding self-reactivity and thus a greater likelihood of becoming CD5^lo^.

To investigate the link between thymic selection efficiency and self-reactivity, as read out by CD5 expression level, we compared the coDP and coSP TCRβ sequences to our previously identified coCD5^lo^ and coCD5^hi^ sequences. Notably, the coDP subsets were slightly more similar to coCD5^lo^ samples, whereas the coSP subsets had a substantially greater overlap in aa sequence with coCD5^hi^ samples (**Figure 6D**). Because all SP sequences must have passed through the DP stage first, the coSP sequences are likely enriched for those that were not captured in the DP stage as a result of them passing through this stage much more quickly. The larger overlap of coSP sequences with coCD5^hi^ samples indicates that their more rapid positive selection may be partly due to higher self-pMHC binding strength, in line with recent experimental results.^45^ In summary, our findings therefore suggest that CDR3β sequences with longer VDJ junctions and acidic aa are less likely to be positively selected and become SP thymocytes, while CDR3β sequences with short VDJ junctions transition from DP to SP thymocytes more quickly and are more likely to become CD5^hi^ cells (**Figure 6E**).

### ML-determined CD5^hi^ propensity scores are predictive of T cell fate differences

Thus far, we established key TCRβ sequence features underlying ML-detected differences in self-reactivity among naïve CD4^+^ T cells. We were able to show, by comparison with TCRs we sequenced from developing thymocytes, that T cells with greater self-reactivity are more efficiently selected into the mature T cell pool. Next, we wanted to investigate whether we could experimentally test ML-generated self-reactivity predictions of TCRβ sequences that were not found in our dataset. To do so, we determined CD5^hi^ propensity scores for six previously investigated TCRβ sequences from CD4^+^ T cells^46^ and determined the two sequences with the lowest and highest propensity scores (**Figure 7A** and **Supplementary Figure S2A**). To ascertain the relative self-reactivity of these two TCRβ chains without fixing the TCRα chain, we generated TCR retrogenic mice expressing the 8-DN (lowest propensity) and 8-24 (highest propensity) TCRβ sequences and measured the CD5 levels on naïve CD4^+^ T cells expressing the TCRβ sequence of interest (**Figure 7B**,**C**, gating strategy shown in **Supplementary Figure S2B**). As might be expected, given that both 8-DN and 8-24 were sequenced from CD4^+^ T cells, the T cell populations in both groups of TCR retrogenics showed a CD4-skewed ratio of CD4^+^ to CD8^+^ T cells, albeit to a greater extent for the 8-DN sequence (**Supplementary Figure S2C**). Importantly, consistent with the ML predictions, CD5 levels on the 8-DN cells were significantly lower than on the 8-24 cells (**Figure 7C**). This confirmed our ML result that the CD5 expression level on naïve CD4^+^ T cells, and thereby relative self-reactivity, could be predicted without knowing the TCRα chain sequence for certain TCRβ chains.

**Figure 7:**
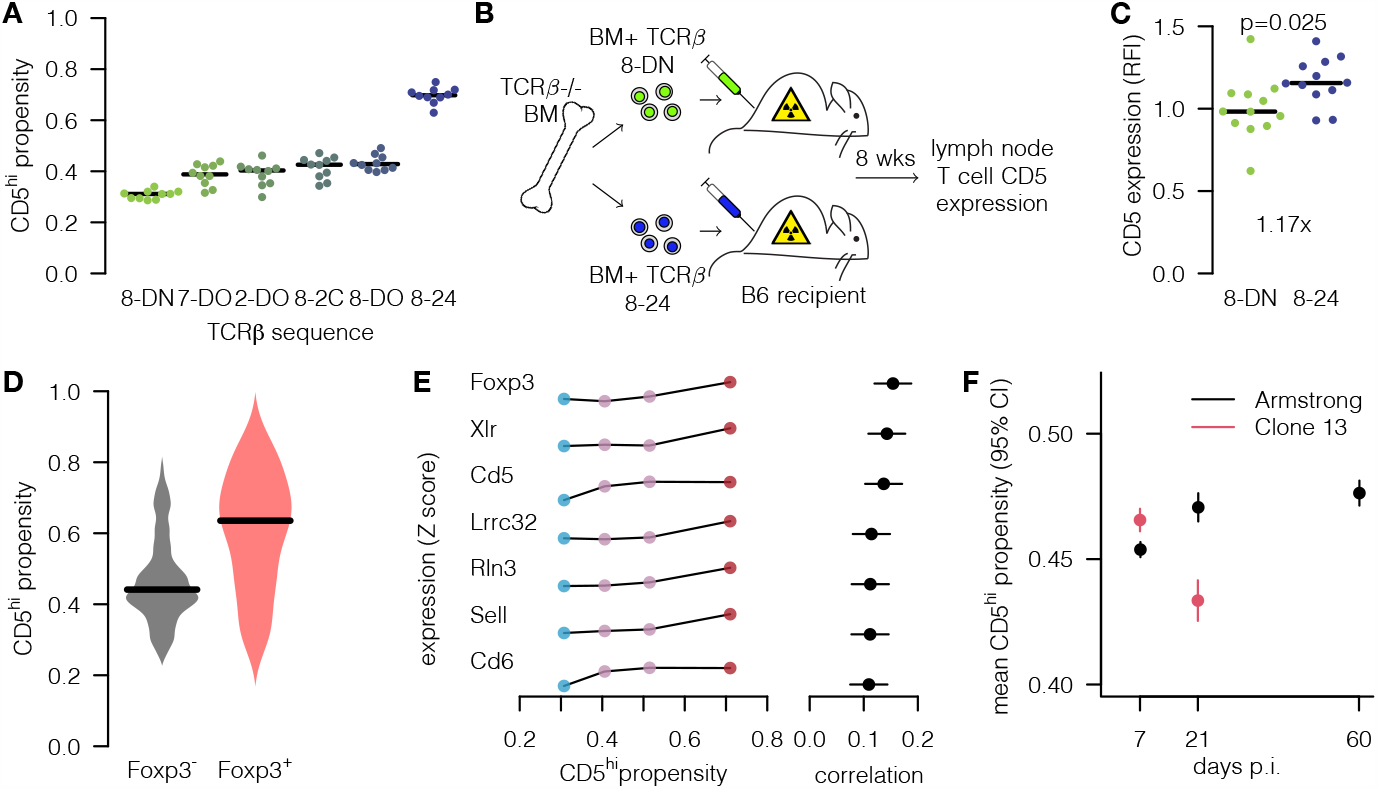
ML-determined self-reactivity scores predict T cell fate differences. **(A)** CD5^hi^ propensity scores assigned by the 10 individual neural networks in the ML ensemble for six CDR3β sequences^46^ that were not part of the initial training or testing data sets. **(B**,**C)** TCR retrogenic mice were generated by transducing TCRβ^-/-^ bone marrow progenitor cells with a retrovirus encoding either the 8-DN or the 8-24 TCRβ sequence (B), and CD5 surface expression levels measured on peripheral naïve CD4^+^ T cells (GFP^+^ to identify cells expressing the transduced TCRβ chain, see also **Supplementary** Figure S2) 6-8 weeks later (C). **(D)** CD5^hi^ propensity score distributions for CDR3β sequences from a published single-cell datasect,^49^ comparing Foxp3^-^ versus Foxp3^+^ CD4^+^ T cells. **(E)** Correlation of ML-determined CD5^hi^ propensity score with gene expression in dataset from (D). Blue and red points indicate the top and bottom 15% of the CD5^hi^ propensity score distribution, with the purple points showing the 15–50% and 50–85% ranges. **(F)** CD5^hi^ propensity scores for LCMV-specific CD4^+^ T cells, comparing the Armstrong strain (acute) and Clone 13 (chronic) strain.^54^

As an additional test of the ML-assigned self-reactivity propensity scores, we next asked whether, even though the ML algorithm was trained only on TCRβ sequences from conventional CD4^+^ T cells, we could extend its predictions to regulatory CD4^+^ T cells (Tregs). Tregs are derived from thymocytes that receive stronger TCR signals during thymic selection, and have, on average, a higher cell-intrinsic self-ligand binding strength than conventional CD4^+^ T cells.^13,47,48^ Therefore, the mean CD5^hi^ propensity score of Treg CDR3β sequences would be expected to be greater than that of conventional CD4^+^ T cells. We analyzed available single-cell RNA-sequencing data of LCMV-specific CD4^+^ T cells where the CDR3β region was also sequenced,^49^ and we used Foxp3 transcript expression to identify Treg cells. In line with previous data and recent comparisons of TCR sequences between conventional and regulatory CD4^+^ T cells,^50^ we found that the Foxp3-expressing T cells had a substantially higher mean CD5^hi^ propensity score (0.61) than Foxp3-negative CD4^+^ T cells (0.47, **Figure 7D**). Using the same dataset, directly correlating CD5^hi^ propensity scores with the expression levels of all detected genes revealed further genes that aligned with the propensity scores, including *Sell* and *Cd6* (**Figure 7E**). Thus, despite being trained only on TCRβ sequences from conventional CD4^+^ T cells, we showed that the ML-identified self-reactivity patterns extended to TCRβ sequences from other CD4^+^ T cell populations and the ML model predicted Treg to have greater self-reactivity based on the TCRβ sequences alone.

Lastly, we applied our ML-generated CD5^hi^ propensity scores to test whether there might be important differences in the self-reactivity of TCR sequences participating in the control of acute versus chronic infections, as had been suggested.^45^,51 Self-pMHC reactivity has been shown to correlate with greater foreign pMHC binding strength in CD4^+^ T cells,^14^ and T cells with greater pMHC reactivity are more prone to exhaustion during chronic antigen stimulation than their low pMHC affinity counterparts.^52,53^ Thus, we investigated the hypothesis that TCRs sequenced from antigen-specific CD4^+^ T cells isolated from a chronic infection setting have a lower self-reactivity than those from an acute infection. We used published single-cell TCRβ sequencing data from CD4^+^ T cells specific for LCMV isolated from mice infected with LCMV-Armstrong, which leads to an acute infection that is cleared in roughly a week, versus LCMV clone-13, which leads to a chronic infection of more than 40 days.^54^ We found that at the peak of the antigen-specific CD4^+^ T cell response (day 7), the CD5^hi^ propensity scores of antigen-specific CD4^+^ T cells identified by tetramer staining in both infections were comparable. In contrast, CD5^hi^ propensity scores were decreased during chronic infection (day 21) compared to antigen-specific memory CD4^+^ T cells sequenced at the same time point post LCMV-Armstrong clearance (**Figure 7F**). Of note, antigen-specific memory CD4^+^ T cells sequenced on day 60 after acute LCMV infection had the highest predicted CD5^hi^ propensities, a finding that is consistent with data suggesting that CD5^hi^ CD4^+^ T cells contribute disproportionately to the memory T cell compartment.^14^

In summary, we applied the ML model trained on our extensive dataset of murine naïve CD4^+^ T cells to predict cell-intrinsic self-pMHC binding strength to various scenarios not covered by the training data, testing key hypotheses using published TCR sequence data. Our results demonstrated that the ML-generated self-reactivity predictions can provide useful TCR sequence-level information on T cell fate in different contexts.

## Discussion

TCR repertoire data is inherently challenging to probe for patterns related to T cell fate. Given the extraordinary number of possible sequences that can be made, many – perhaps most – sequences are unique within an individual. This means that common approaches, including differential gene expression (DGE) analysis, which rely on observing the same sequence repeatedly across individuals to reach statistical significance, are not well suited for TCR sequence data. Indeed, much of our understanding of which TCR sequences predominate in a T cell repertoire has come from quantifying generation probabilities, showing that public sequences which reoccur across individuals are the product of convergent recombination (many different recombination events can give rise to the same nucleotide sequence) compared to private TCR sequences.^30^ However, this entirely stochastic view of TCR repertoire generation has been difficult to reconcile with our understanding of thymic selection processes and MHC restriction, whereby only T cells with a sufficient ability to interact with the self-peptides presented by the MHC alleles of an individual are positively selected.^55^ Moreover, it is well-documented that the signal strength obtained by thymocytes has important ramifications for cell fate: T cells with greater self-reactivity are more likely to develop into Treg cells or be removed from the repertoire by negative selection,^47,56^ and positive selection has been shown to be less efficient for T cells with low self-reactivity.^45^ Importantly, at the whole-TCR repertoire level, biases in representation due to self-reactivity among naïve T cells have been difficult to discern, and to what extent the TCR sequence can predict self-reactivity and therefore cell fate outcomes has been an open question.

Here we show that traditional analysis methods of TCRβ sequencing data generated from naïve CD4^+^ T cells sorted into populations with high and low self-reactivity, based on CD5 surface level expression, provided only limited ability to detect differences in sequence features. Therefore, we turned to ML to identify subsets of the sequence datasets that were more specific to each population. ML is increasingly being applied to immunology, often for classification tasks such as linking TCR sequences to known epitopes. While the ML model we implemented is technically a classifier, we did not use it as such. Instead, we used our ML model as a filter to identify relevant TCRβ sequence subsets to focus our analysis on. In that sense, our ML model is more comparable to a DGE analysis, but with the key difference that it also learns from the many sequences observed only once. Indeed, this general approach to use an ML classifier in lieu of a DGE analysis for TCR sequencing data will likely be applicable to other studies as well, given that the presence of many unique clones is a hallmark of TCR repertoires.

The use of an ML algorithm revealed several interesting TCR sequence features that correlate with self-reactivity. For the first time, we provide evidence that, in addition to the known impact of non-templated nt-additions in reducing the TCR clone size, positive selection efficiency, and cross-reactivity,^57,58^ TCRs with more n-nucleotides tended to have reduced self-reactivity, as had been previously hypothesized.^10^ We and others have shown that the expression of TdT at the gene-level is substantially reduced (>10 fold) in higher-affinity T cells in both naïve CD4^+^ and CD8^+^ T cells, in humans and mice, although this had not been previously observed to be reflected by n-nucleotide additions at the TCR sequence level.^17,21,22,37^ Together with the relationship between *Dntt* expression and self-reactivity, our findings raise the intriguing possibility that there could be a trade-off between TCRβ repertoire diversity – which is increased by *Dntt* activity – and recognition strength – which is decreased by *Dntt* activity. It will be interesting to test this idea directly in a system where *Dntt* expression levels can be raised or lowered experimentally, TCRs sequenced, and subsequent cell fates and repertoire diversity studied. In being able to stratify the TCR sequences in our dataset based on the confidence with which the ML algorithm assigned it to high- or low-self reactivity subsets, we also unraveled specific features of TCR aa sequences that were associated with strong self-reactivity, including an enrichment of hydrophobic and basic aa, while other aa were associated with weaker self-reactivity, such as acidic aa. The largest differential aa usage patterns were seen in the middle positions of the CDR3β, lending support to the idea that these positions could be the most important ones for determining specificity,^59^ and in agreement with prior noted patterns based on the study of specific TCRs.^60^ Since use of ML algorithms comes with a risk of “overfitting” the data, and interpreting ML classifier results is not straightforward, we confirmed the importance of these sequence features by analyzing an independent dataset in an ML-independent manner, pre-publishing our analysis plan for transparency and greater confidence in the statistical analysis.

Overall, we made the surprising finding that the CDR3β sequence alone can, in some instances, determine CD4^+^ T cell fate – irrespective of the α chain. We showed that TCR sequences from CD5^lo^ TCRs are slightly overrepresented among pre-selection DP thymocytes, implying that they might take longer to accrue sufficiently many productive TCR signals to be positively selected, as experimental data had suggested.^45^ Conversely, TCR sequences from CD5^hi^ TCRs were underrepresented among pre-selection DP thymocytes, suggesting they pass rapidly through this stage as they more quickly acquire the necessary TCR signal. This would be in line with the CD5 expression distribution skew previously identified,^15^ reflecting less variability in the selection trajectories of higher-affinity cells. Moreover, we corroborated the ML-based self-reactivity predictions using TCR retrogenic mice where we fixed two TCRβ sequences, but where the α chain was left to vary, and we experimentally observed the predicted differences in relative CD5 levels. In extending the application of the ML algorithm to publicly available datasets, we found Tregs’ TCRs have much higher CD5^hi^ propensity scores, consistent with previous studies.^15^ This finding also suggests that TCRs from Treg follow similar rules regarding TCR self-reactivity. Indeed, a recent meta-analysis of all published Treg versus naive CD4^+^ T cell TCRβ sequences used statistical models to show that self-reactivity is to some extent a function of the TCR sequence and described similar aa usage patterns as we do here.^50^ Lastly, we applied the ML model to a TCR sequencing data set from T cells responding to acute versus chronic LCMV infection, showing that the mean CD5^hi^ propensity score drops during chronic infection, potentially reflecting preferential exhaustion of strongly self-reactive cells.^45,51^

Of interest, many TCR sequences in our dataset could not be confidently mapped to their CD5 status. There are two possible explanations for this. First, the α chain sequence may be required to enable accurate CD5 level prediction for those sequences. Second, the CD5-level might generally be a result of not only the TCR sequence but also of stochastic interactions with self in the thymus, leading to a non-deterministic relation between TCRβ sequence and CD5 expression. These two possibilities remain to be tested using datasets where both α and β chains of individual T cells are known, and CD5 levels are determined at the protein level.

In summary, our work has allowed us to define basic features of the architecture of a TCR repertoire architecture. We have established that the TCRβ sequence can, by itself, be predictive of the fate and function of naïve CD4^+^ T cells. Because the TCR is what ultimately defines the identity of a T cell, further unravelling the principles linking TCR sequence to TCR function will fundamentally advance our understanding of T cell biology.

## Acknowledgements

We would like to thank P. D’Arcy, G. Perreault, and animal facility staff at McGill for their excellent care of our animal colony, Ann Feeney (Scripps) for sharing TdT^-/-^ mice, Thierry Mallevaey for providing the TCRβ constructs used for the retrogenic experiments, Julien Leconte and Camille Stegen at the McGill University Cell Vision Core Facility for performing all cell sorts, and P. Artusa for contributions to T cell sorting. DR was funded by a Frederick Banting and Charles Best Canada Graduate Scholarships Doctoral Award (CIHR CGS-D) and a Tomlinson Doctoral Fellowship (McGill University). EMG is the recipient of a Natural Sciences and Engineering Research Council of Canada Graduate Scholarship Doctoral award (CGS D). HJM is a Fonds de recherche du Quebec-Sante Senior Research Scholar. JNM holds a Canada Research Chair for immune cell dynamics. This research was supported by a NSERC Discovery Grant (2016-03808) and a McGill start-up fund to JNM, a CIHR (PJT-168862) grant jointly to JNM and HJM, and an NWO Vidi grant (VI.Vidi.192.084) to JT. JW was funded by the German Research Foundation (DFG) within the framework of the Schleswig-Holstein Excellence Cluster I&I (EXC 306, Inflammation at Interfaces, project XTP4), the graduate schools GRK 1727/2 and the TR-SFB654 project C4 at the University of Lübeck.

## Author contributions

JNM and JT conceived, designed and supervised the project and research, with key input from HJM and IW. FB and JT analyzed the TCR sequencing data. FB developed the machine learning algorithm with input from SS under supervision by JT. Under the supervision of JW, KK, AF and RP performed all TCR-seq on cells sorted by DR and JNM. EMG performed the TCR retrogenic mouse experiments designed and supervised by HJM. The manuscript was written by JNM, JT, DR and FB with feedback from all other authors.

## Materials and Methods

### Mice

C57BL/6, RAG2-GFP,^61^ and TCRβ^-/-62^ breeders were obtained from Jackson Laboratories. TdT^-/-^ mice^6^ were kindly shared by A. Feeney (Scripps) and bred in-house. All mice were on a C57Bl/6 background and used for experiments at 6-12 weeks of age, with both males and females used in experiments, except for TCR sequencing cell sorts where only female mice aged 8 weeks were used. Animal housing, care, and research were in accordance with the Guide for the Care and Use of Laboratory Animals and all procedures performed were approved by the McGill University Animal Care Committee. For the retrogenic mice, the animal protocol was approved by the Animal Care Committee and experiments performed at the Maisonneuve-Rosemont Hospital Research Centre.

### Thymocyte and lymphocyte isolation

For experiments with peripheral naïve CD4^+^ T cells, spleen and peripheral lymph nodes (inguinal, axillary, brachial, superficial cervical, and mesenteric) were harvested and passed through a 70μm filter with 1% RPMI (1% FBS, 1% L-glutamine, and 1% pen/strep). ACK lysis buffer (Life Technologies) was added for 3 minutes, and samples were re-filtered and resuspended in 1% RPMI. For experiments with thymocytes, each thymus was harvested and passed through a 70μm filter with 1% RPMI (1% FBS, 1% L-glutamine, and 1% pen/strep). Cell counts were determined by diluting a single-cell suspension 1:10 in Trypan Blue (ThermoFisher Scientific) and manually counting live single cells (trypan blue-negative).

### Flow cytometry

Cells were incubated in Fixable Viability Dye (AF780, eBioscience) diluted in PBS for 20 minutes at 4°C. Extracellular antibodies were diluted in FACS buffer (2% FBS and 5mM EDTA in PBS) with added Fc Block (1:100, eBioscience) and incubated for 30 minutes at 4°C. Samples requiring intracellular staining were subsequently incubated in FoxP3 Transcription Factor Fixation/Permeabilization Concentrate and Diluent (Life Technologies) for 30 minutes at 4°C. Intracellular antibodies were diluted in permeabilization wash buffer and incubated for 60 minutes at 4°C. Directly conjugated antibodies used were as follows: TCRβ (H57-597), CD3 (145-2C11), CD4 (RM4.5), CD8 (53-6.7), CD25 (PC61.5), CD44 (IM7), CD62L (MEL-14), CD5 (53-7.3), FoxP3 (FJK-16s), TdT (19-3), Ly6C (HK1.4), B220 (RA3-6B2), CD11b (M1/70), CD11c (N418), F4/80 (T45-2342), NK1.1 (PK136). For all flow cytometry experiments of naïve CD4^+^ T cells, Tregs were excluded by intracellular staining for FoxP3, and cells were acquired using an LSRFortessa (BD Bioscience) and analyzed with FlowJo software (BD Bioscience).

### Cell sorts

Thymocytes and lymphocytes from C57Bl/6 or C57Bl/6.SJL (CD45.1^+^) congenic mice were isolated in single cell suspension as described. Total isolated thymocytes were directly stained for sorting. Total lymphocytes were magnetically enriched for CD4^+^ T cells (Stemcell EasySep mouse CD4^+^ T cell enrichment kit). Cells were then incubated in Fixable Viability Dye and subsequently stained with surface antibodies for 1 hour at 4°C. Sorts were performed on either a FACS Aria Fusion, Aria III, or Aria II SORP (BD Bioscience). All cell populations were sorted to >90% purity for bulk populations.

*Sorts for TCR-seq:* Peripheral naïve CD4^+^ T cells were sorted on live, TCRβ^+^, CD4^+^, CD8^-^, CD25^-^ (to exclude Tregs), CD44^-^, CD62L^+^, and 25% CD5^lo^ or CD5^hi^ (**Supplementary Figure S1C**,**D**). Thymocytes were sorted on live and TCRβlo, CD5^lo^, CD4+ and CD8+ for pre-selection DP cells; and TCRβ^hi^, CD8^-^, CD4^+^ and CD25^-^ for CD4^+^ SP cells (**Supplementary Figure S1E,F**). Of note, we previously confirmed that sorted peripheral CD5^lo^ or CD5^hi^ naïve CD4^+^ T cells using this sort strategy did not contain Tregs post-sorting by measuring Foxp3 expression using qPCR.^21^ *Sorts for quantitative RT-PCR:* Thymocytes were sorted on lineage negative (B220, CD11b, CD11c, F4/80, Ly6C, and NK1.1) and TCRβ^lo^, CD8^-^, and CD4^-^ for DN; TCRβ^lo^, CD8^+^, CD4^+^ for DP; and TCRβ^hi^, CD8^-^, CD4^+^ for CD4^+^ SP. Peripheral naïve CD4^+^ were sorted on lineage negative, CD25^-^, TCRβ^+^, CD4^+^, CD8^-^, RAG2^-^GFP^-^, CD44^-^, and 20% CD5^lo^, CD5^mid^, or CD5^hi^.

### RNA extraction and quantitative real-time RT-PCR

RNA from sorted DN, DP, and SP thymocytes and 20% CD5^lo^, CD5^mid^, or CD5^hi^ naive CD4^+^ T cells was extracted using RNAqueousTM-Micro Total RNA Isolation Kit (Life Technologies) and cDNA converted using High-Capacity cDNA Reverse Transcription Kit (Life Technologies). qPCR analysis was performed with TaqManTM Gene Expression Master Mix (Life Technologies) and TaqManTM Gene Expression Assay (FAM, *Dntt*, Mm00493500_m1, Life Technologies). Average Ct values across technical duplicates were determined for *Dntt* and fold change was calculated as Log2-transformed 2ΔΔCt values relative to the expression of the housekeeping gene Gapdh.

### TCRβ sequencing

RNA from sorted populations was extracted using innuPREP RNA Mini Kit (Analytik Jena, Hildesheim, Germany). cDNA synthesis, amplification of TCRβ-chain transcripts and library preparation was performed with the arm-PCR (amplicon-rescued multiplex PCR) technology (iRepertoire Inc. Huntsville, USA) using the Qiagen OneStep RT-PCR Kit and Qiagen Multiplex PCR Kit (both Qiagen) according to the manufacturer’s protocols. PCR products were run on a 2% agarose gel and purified using QIAquick Gel Extraction Kit (Qiagen, Hilden, Germany). The obtained TCRβ libraries were quantified using the PerfeCTa-NGS-Quantification Kit according to the manufacturer’s protocol (Quantabio Inc, Beverly, USA) and sequenced using the Illumina MiSeq Reagent Kit v2 300-cycle (150 paired-end read; Illumina) and the MiSeq system (Illumina Inc. San Diego, USA). We merged the paired-end reads using PEaR,^63^ and mapped the CDR3β region, clustered clonotypes, and corrected sequencing errors such as removal of nonfunctional CDR3β sequences using the Recover TCR pipeline.^64^

### Generation of TCRβ retrogenic mice

TCRβ constructs (8-24 or 8-DN) containing a GFP reporter gene were generated as previously described^46,65,66^ and kindly provided by Thierry Mallevaey (University of Toronto). 293T cells (ATCC) were transfected with the above retroviral plasmids encoding the 8-24 or 8-DN TCRβ transgene and the pCL-Eco packaging vector (Addgene) using Lipofectamine 2000 (Life Technologies) according to manufacturer instructions.^67^ Retroviral supernatant was collected 48hr and 72hr after transfection and either used immediately or stored at 4°C for up to 24 hours for transduction of bone marrow cells. Retrogenic TCR mice were made as described by Holst *et al*.^68^ with some modifications. TCRβ^-/-^ mice were injected intraperitonially with 5-fluorouracil (15 mg/g weight; Accord Healthcare). Four days later, bone marrow cells were harvested from the femurs, tibiae, and ilia and stimulated overnight in DMEM (Wisent) containing 20% FBS (Thermo Fisher Scientific), 10-5 M β-Mercaptoethanol (Sigma-Aldrich), 50 IU/mL-50μg/mL penicillin-streptomycin (Wisent), 20 ng/mL IL-3 (Biolegend), 50 ng/mL IL-6 (Biolegend), and 50 ng/mL stem cell factor (Biolegend). Bone marrow cells were transduced with retroviral supernatants supplemented with Polybrene (6 µg/mL, Sigma) by spinfection. Briefly, the fresh retroviral supernatant was added in a 1:1 volume ratio to the bone marrow cells in a 6-well plate and spun at 1000g for 1hr at room temperature. The spinfection was repeated 24h later with 1 mL of fresh viral supernatant, and the cells were subsequently maintained in the incubator at 37°C for 4-10 hours. The bone marrow cells were then spun at 225g for 10min at 4°C and resuspended at a concentration of 10^7^ cells/mL. 300µL of the cell suspension was injected intravenously into lethally irradiated (10gy) C57BL/6 CD45.1^+^ recipient mice. Spleens from retrogenic mice were analyzed by flow cytometry 6-8 weeks after reconstitution. Mice with <2.5% GFP^+^ T cells of the live non-B cell population were not included in CD5 expression level comparisons.

### Machine learning

To identify systematic differences between CD5^lo^ and CD5^hi^ CDR3β sequences, an artificial neural network (ANN) was trained to predict whether a sequence, V gene and J gene comes from a CD5^lo^ or CD5^hi^ cell. The sequences that the network was able to correctly identify confidently as CD5^lo^ or CD5^hi^ were extracted and compared on their V-J distance, V-gene, and J-gene usage. The SP versus DP, WT vs TdT^-/-^ (positive control) and WT vs WT (negative control) ANNs were trained and evaluated according to the same protocol described below for the CD5^lo^ vs CD5^hi^ case.

#### Preprocessing

For training an ANN to distinguish CD5^lo^ and CD5^hi^ CDR3β sequences we used data from 19 mice (**Figure 1**), which we preprocessed as follows. Firstly, sequences longer than 64 nucleotides (which are likely mapping errors), as well as sequences with only one mapped read (which have the highest risk of containing sequencing errors), were removed from the analysis. As the goal of this study was to find unique features for CD5^lo^ or CD5^hi^ cells, sequences shared between CD5^lo^ and CD5^hi^ were removed from the dataset during training. Duplicates were defined as having the same sequences, V gene, and J gene, and that were present in both CD5^lo^ or CD5^hi^ cells. Second, the data was preprocessed to match the input shape required for the ANN. The TCR sequences were first padded to an equal length of 64, the maximum length for sequences still present in the dataset. As in a prior study,^41^ the TCRβ sequence was split into two equal parts, and padding was appended in the middle of the two parts. In the case of an odd length sequence, one extra nucleotide was added to the left part. The padded sequences were then one-hot encoded to a shape of 64 by 5, corresponding to the final length of the sequences and the five categories present: the four nucleotides and the padding. The V genes and J genes were one-hot encoded as well. Since there are in total 22 different V genes and 11 different J genes, the final length of a one-hot encoded V gene and a one-hot encoded J gene are 22 and 11 respectively. The output was encoded as zero or one: zero corresponding to CD5^lo^ and one corresponding to CD5^hi^. After encoding, the data were randomly split into a training set and validation set, with an 80/20 ratio. After the train/test split, sequences were removed from the test set if their aa sequence corresponded to the aa sequence in the training set, to make the test set unique with respect to the aa sequence. To prevent the model from overfitting, the classes were balanced by presenting an equal amount of CD5^lo^ or CD5^hi^ sequences to the model during training. Lastly, the data was randomly shuffled before training.

#### Model architecture

The model consists of three input layers (**Figure 3B**): one for the sequences, one for the V genes, and one for the J genes. Each input was then fed through four fully connected layers with 64 units and Rectified Linear Unit (ReLU) as activation function before they were concatenated. After concatenation, the concatenated features were again fed through three fully connected layers with 64 units and ReLU activation. Lastly, the output layer consisted out of one node and sigmoid activation function. A prediction of 0 represented CD5^lo^ and a prediction of 1 represented CD5^hi^.

#### Training

The model was trained with a batch size of 512. Adam^69^ was used as optimizer with a learning rate of 10^−4^. Binary cross-entropy was used as a loss function. To prevent overfitting during training and improve the models’ generalization, early stopping was used. The validation loss was monitored as a performance measure, and training was stopped when the validation loss did not decrease for 6 epochs.

### Analysis

After training the model and validating that the model was able to truly learn differences between the two classes on the test data, the trained model was used to determine the CD5 propensity for all CD5^lo^ and CD5^hi^ sequences. Per mouse, the 15% CD5^lo^ sequences with the lowest CD5^hi^ propensity and the 15% CD5^hi^ sequences with the highest CD5^hi^ propensity were selected. These were used as CD5^lo^-characteristic sequences (coCD5^lo^) and CD5^hi^-characteristic sequences (coCD5^hi^) for further analysis.

### Statistical analysis

All statistical analyses were performed within the R platform for statistical computing. All analysis scripts will be made available at this paper’s GitHub repository at github.com/jtextor/tcr-self-reactivity. Briefly, P values were computed using paired (**Figure 2E**,**Figure 4B**) or unpaired (**Figure 7C**) t tests. Amino acid enrichment plots (**Figure 4F**,**Figure 6C**) were made by scaling the height of each letter at each position according to the standardized residual of a chi-square test that compared the amino acid distributions between the two groups at that position. Differential gene expression analysis (**Figure 2G**,**H**) was conducted using the edgeR Biocondoctor package.^70^ Nt nucleotides (**Figure 2G**,**H**) were estimated by aligning the D1 and D2 segment sequences of the murine TCRβ chain ot the junction sequence, determining the longest alignment, and counting the number of nucleotides in the junction that were not part of this alignment, the V segment, or the J segment. Validation analyses (**Figure 5)** were conducted as described in our data analysis plan.^42^ Briefly, usage of V and J genes of interest was encoded as a binary variable. To measure junction acidity and hydrophobicity, the percentage of acidic amino acids (glutamic or aspartic acid) or hydrophobic amino acids (leucine, isoleucine, valine or phenylalanine) contained in the junction sequence – except the initial 3 and final 2 positions – was determined. Together with V-J distance, this resulted in 11 features per sequence. For feature-by-feature analyses, we grouped CD5^lo^- and CD5^hi^ associated gene segments by taking a logical “or” of the corresponding binary variables.

## Data Availability

Raw TCR sequencing data, as well as processed data from the RTCR pipeline, have been deposited on GEO (accession number: GSE221703; BioProject ID: PRJNA915397). Additionally, the validation dataset including CD5 propensity predictions and data analysis code (**Figure 6**) have been deposited on Zenodo.^42^

## Code Availability

A Python implementation of the CD5 propensity prediction algorithm is available at this paper’s GitHub repository at github.com/jtextor/tcr-self-reactivity. We also provide an web browser based implementation of the algorithm at https://computational-immunology.org/cd5-prediction/.

## Supplementary Figures

**Supplementary Figure S1:**
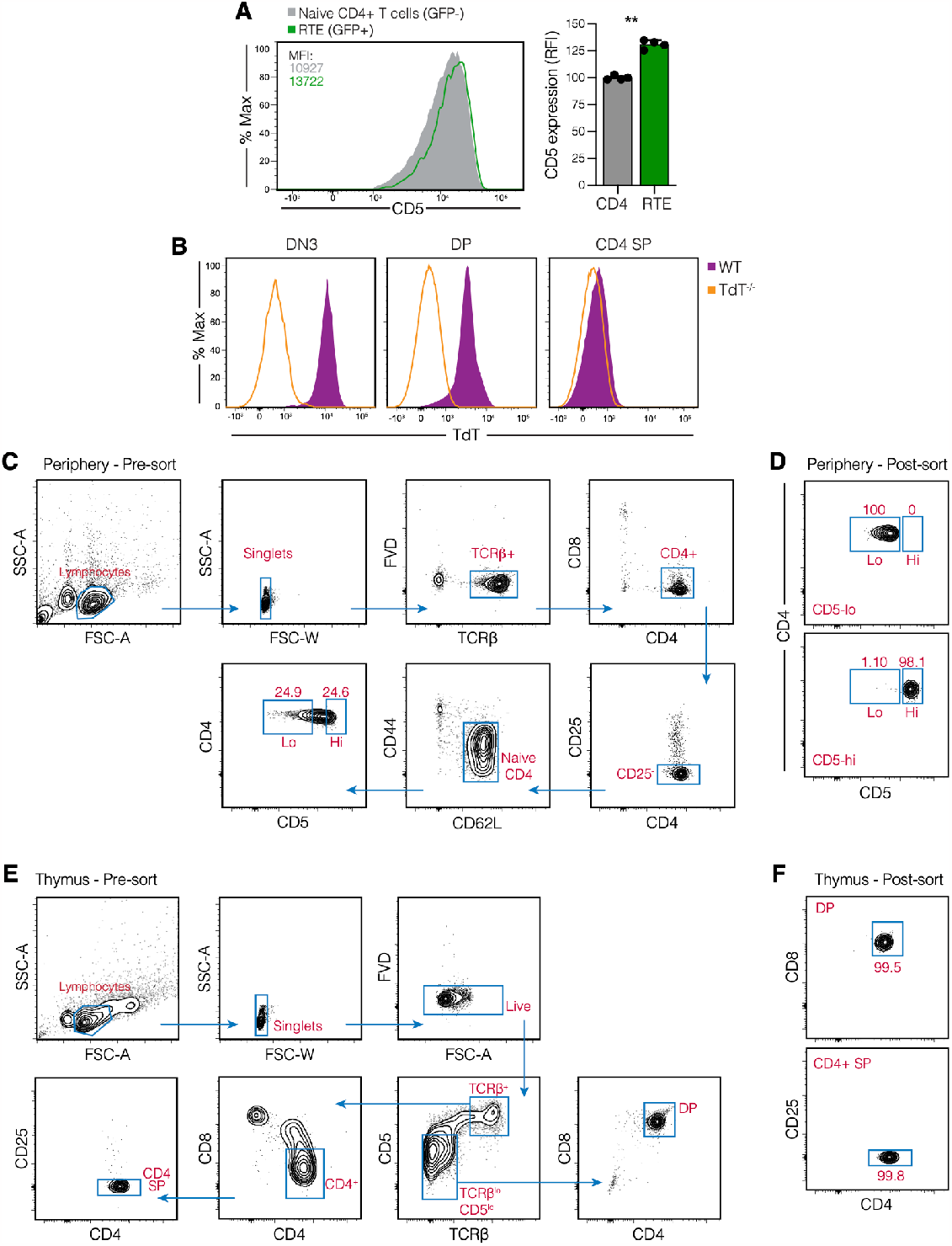
Greater *Dntt* expression in CD5^lo^ CD4^+^ T cells is not explained by contribution of recent thymic emigrants. **(A)** Representative flow cytometry histogram (left) and summary data (right) of CD5 surface expression levels on recent thymic emigrants (RTE) identified using a Rag-GFP reporter, compared to non-RTE naïve CD4^+^ T cells. **(B)** TdT protein expression measured by flow cytometry on thymocytes in the double negative (DN3), double positive (DP) and CD4^+^ single positive (SP) stage of T cell development in wild type (WT) and TdT^-/-^ mice. **(C-F)** Gating strategy for peripheral T cell populations pre- (C), and post- (D) sort, as well as thymic T cell populations pre- (E) and post- (F) sort for TCRβ sequencing. Numbers in plots indicate percent cells in gate; FVD, fixable viability dye.

**Supplementary Figure S2:**
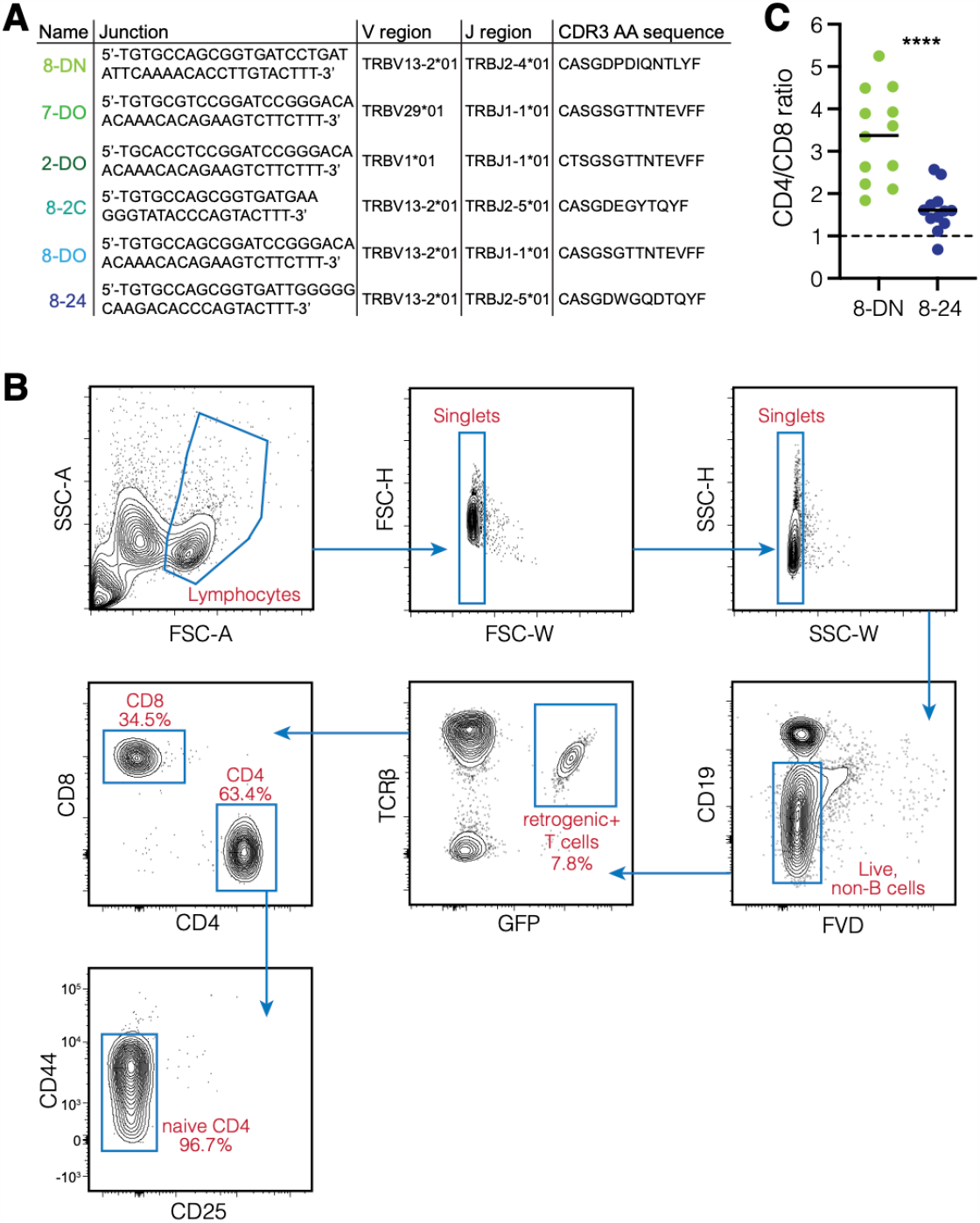
Generation of TCRβ retrogenic mice. **(A)** Junction nucleotide sequences, V/J gene usage, and amino acid CDR3β sequences for the TCRβ sequences for which CD5^hi^ propensity scores were determined. **(B)** Gating strategy used for assessing CD5 surface expression levels on naïve CD4^+^ T cells generated by bone marrow progenitor cell transduction with the 8-DN or the 8-24 TCRβ sequence. Retrogenic T cells were identified by GFP expression. FVD, fixable viability dye. **(C)** CD4^+^ to CD8^+^ T cell ratios of 8-DN and 8-24 retrogenic T cells. Data is summarized from 4 independent experiments; each data point is from an individual mouse (N=12).

